# Reply to: Contribution of carbon inputs to soil carbon accumulation cannot be neglected

**DOI:** 10.1101/2023.08.20.552557

**Authors:** Feng Tao, Benjamin Z. Houlton, Serita D. Frey, Johannes Lehmann, Stefano Manzoni, Yuanyuan Huang, Lifen Jiang, Umakant Mishra, Bruce A. Hungate, Michael W. I. Schmidt, Markus Reichstein, Nuno Carvalhais, Philippe Ciais, Ying-Ping Wang, Bernhard Ahrens, Gustaf Hugelius, Toby D. Hocking, Xingjie Lu, Zheng Shi, Kostiantyn Viatkin, Ronald Vargas, Yusuf Yigini, Christian Omuto, Ashish A. Malik, Guillermo Peralta, Rosa Cuevas-Corona, Luciano E. Di Paolo, Isabel Luotto, Cuijuan Liao, Yi-Shuang Liang, Vinisa S. Saynes, Xiaomeng Huang, Yiqi Luo

## Abstract

In the accompanying Comment^1^, He et al. argue that the determinant role of microbial carbon use efficiency in global soil organic carbon (SOC) storage shown in Tao et al. (2023)^2^ was overestimated because carbon inputs were neglected in our data analysis while they suggest that our model-based analysis could be biased and model-dependent. Their argument is based on a different choice of independent variables in the data analysis and a sensitivity analysis of two process-based models other than that used in our study. We agree that both carbon inputs and outputs (as mediated by microbial processes) matter when predicting SOC storage – the question is their relative contributions. While we encourage further studies to examine how the evaluation of the relative importance of CUE to global SOC storage may vary with different model structures, He et al.’s claims about Tao et al. (2023) need to be taken as an alternative, unproven hypothesis until empirical data support their specific parameterization. Here we show that an additional literature assessment of global data does not support He et al.’s argument, in contrast to our study, and that further study on this topic is essential.

## Main

The higher explanatory power of carbon input than microbial CUE to SOC storage envisaged by He et al. does not hold at the global scale when more data are considered (Table 1). He et al. proposed that carbon input is potentially more important than microbial CUE by using the net primary productivity (NPP) as carbon input to explain the spatial variation of SOC in 132 datasets in a mixed-effects model. However, the statistical models in Tao et al. (2023) were not applied to evaluate the relative importance of either CUE or NPP to SOC, but to determine whether microbial CUE is positively or negatively correlated with SOC. This issue raised by He et al. might become relevant if NPP obscures the CUE-SOC relationship to the extent of changing its direction. In Supplementary Table 3, Tao et al. (2023) showed that the inclusion of NPP in a mixed-effects model does not influence the positive CUE-SOC correlation. Moreover, NPP may have high explanatory power for SOC across these 132 local sites, but not necessarily at the global scale. We extracted NPP information from a MODIS-based product^3^ and the PRODA-retrieved CUE from 57,267 soil profiles (Extended Data Fig. 1b of Tao et al., 2023) and found that CUE explains more spatial variation of SOC (37%) than NPP (9%) at the global scale (Table 1). Indeed, the notion that NPP is a minor factor in explaining SOC dynamics and spatial variation at both regional and global scales has been well documented in the literature^4-6^.

**Table 1.** Microbial CUE explains more spatial variation of SOC storage than NPP at the global scale. Statistics shown in the table are unstandardized coefficients of relationships between CUE (from data assimilation) or NPP (from remote sensing data) and SOC content in a mixed-effects model. CUE or NPP was set as the fixed effect to predict SOC content. Climate types that soil profiles belong to were set as the random effect. We assumed random intercepts in all regressions. The total observation size *n*_*obs*_ = 56,270, the random effect size n_climate_ = 12.

|  |  | Intercept | CUE or NPP |
| --- | --- | --- | --- |
| *log10(SOC)~CUE + (1\|Climate Type)* |  |  |  |
| **variance explained by mixed model: 37%** |  |  |  |
| Fixed Effects | Estimates | 0.89 | 1.18 |
|  | Std. Error | 0.055 | 0.0096 |
|  | t value | 16.07 | 122.44 |
|  | P | <0.0001 | <0.0001 |
| Random Effects | Standard Deviation | 0.19 | NA |
| *log10(SOC)~NPP + (1\|Climate Type)* |  |  |  |
| **variance explained by mixed model: 9%** |  |  |  |
| Fixed Effects | Estimates | 1.13 | $2.13 \times 10^{-4}$ |
| | Std. Error | 0.065 | $4.61 \times 10^{-6}$ |
|  | t value | 17.54 | 46.32 |
|  | P | <0.0001 | <0.0001 |
| Random Effects | Standard Deviation | 0.05 | NA |

**Fig. 1.**
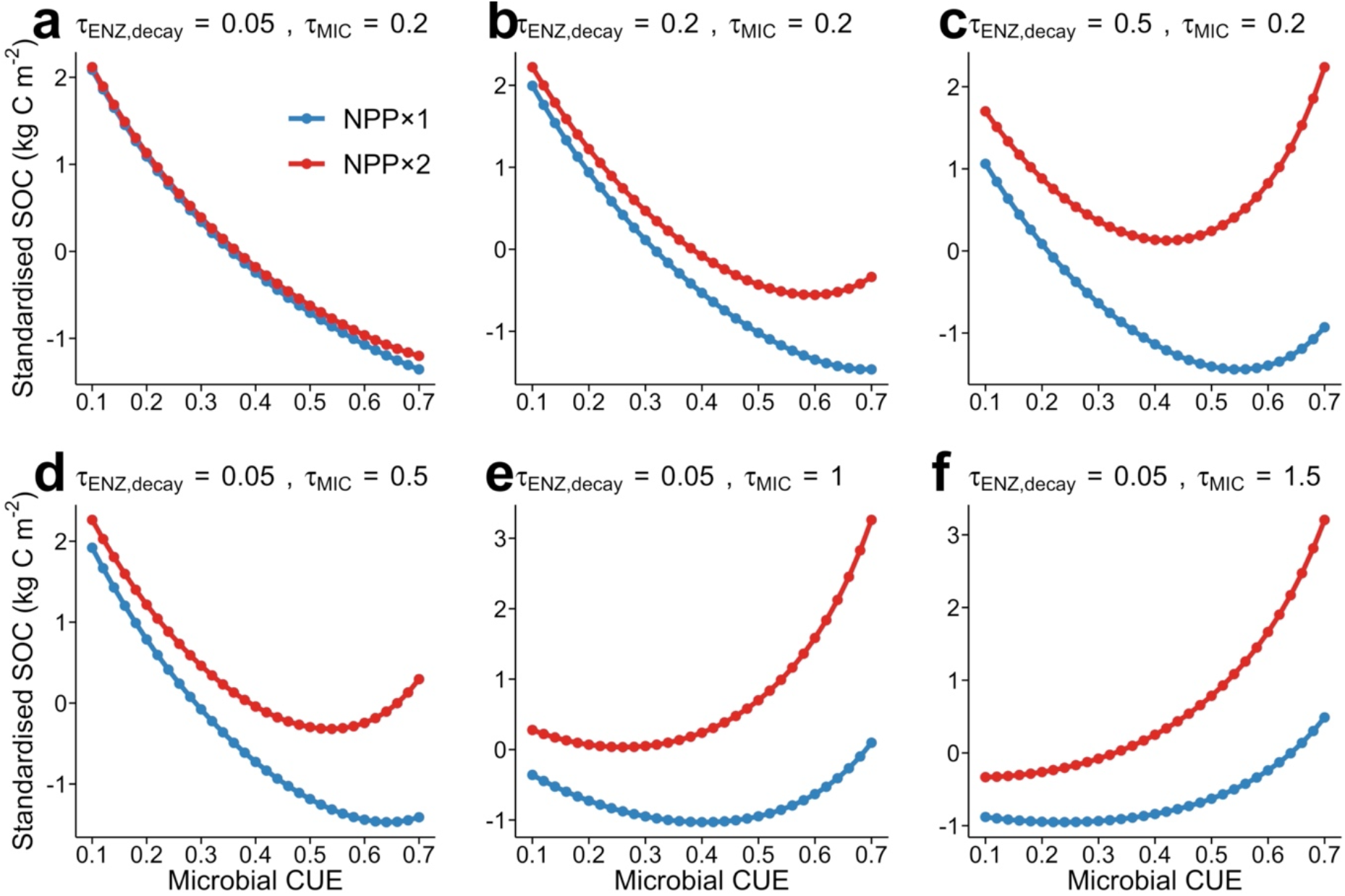
Varying sensitivity of SOC storage to doubled NPP under different combinations of parameter values with the microbial model used in Tao et al. (2023). We chose different values for the turnover time (in years) for enzyme and microbial biomass carbon pools of the microbial model used in Tao et al. (2023) at one site and doubled the magnitude of NPP in model simulation for each set of parameterizations. Different panels in this figure present how the relation of CUE and SOC storage changed with increased NPP. The SOC values were standardized using the Z-score method to be comparable with the results shown in He et al.

Process-based models can theoretically generate an array of patterns and predictions, as correctly argued by He et al. and as well documented in the literature. However, models yield realistic predictions only after they are constrained by observations. He et al. used two models to examine varying sensitivities of SOC storage in response to changes in a parameter, *β*, that represents the density dependence of microbial mortality^7^, arguing that SOC storage could be more sensitive to changes in NPP than in microbial CUE under certain parameterizations. Similarly, the microbial model used in Tao et al. (2023) generates a nested set of sensitivities for SOC storage in response to NPP, but assuming that *β* = 1 – i.e., assuming that mortality is not density-dependent (Fig. 1). SOC storage shows no response to doubled NPP when the turnover time (*τ*) of both enzyme (ENZ) and microbial biomass (MIC) is very short (e.g., *τ*_*ENZ,decay*_ = 0.05 years and *τ*_*MIC*_ = 0.2 years, Fig 1a). Alternatively, SOC storage dramatically increases with doubled NPP when the turnover time of either of these two pools is higher (e.g., *τ*_*ENZ,decay*_ increases from 0.05 years to 0.5 years, Fig. 1a-c, or *τ*_*MIC*_ increases from 0.2 years to 1.5 years, Fig. 1d-f). Nevertheless, microbial CUE emerged from our SOC data assimilation into the model to be more important than NPP to global SOC storage. It is the Bayesian framework used in our study via data assimilation that identified the most probable mechanisms among these diverse mechanisms. We look forward to an analysis where first data assimilation is conducted to estimate the *β* value in order to then demonstrate that the estimated parameter could disprove the fundamental importance of microbial CUE to SOC storage.

Estimates of NPP by different process-based models and data products indeed remain uncertain, as pointed out by He et al. While the uncertainty in NPP might influence CUE-SOC relationships, our analysis showed that variation in NPP values from -10% to +10% of the CLM5 simulated values had much less effect on SOC than microbial CUE (Fig. 4b of Tao et al., 2023). Nevertheless, we greatly appreciate the suggestion to include carbon input as a parameter for optimization. We thus encourage the scientific community to conduct data assimilation studies to constrain all parameters, including mortality- and NPP-related ones, that may influence the CUE-SOC relationships.

## Methods

All the data, statistical methods, and the microbial model have been described in Tao et al. (2023) and can be publicly accessed via https://www.nature.com/articles/s41586-023-06042-3.

## Author contributions

F. T. and Y. L. drafted the reply. All authors contributed to the text and approved the final version.

